## Supplementary figures and images for "HI-FISH: WHOLE BRAIN IN SITU MAPPING OF NEURONAL ACTIVATION IN DROSOPHILA DURING SOCIAL BEHAVIORS AND OPTOGENETIC STIMULATION"

### Figure 1 - Supplement

#### Figure 1 - figure supplement

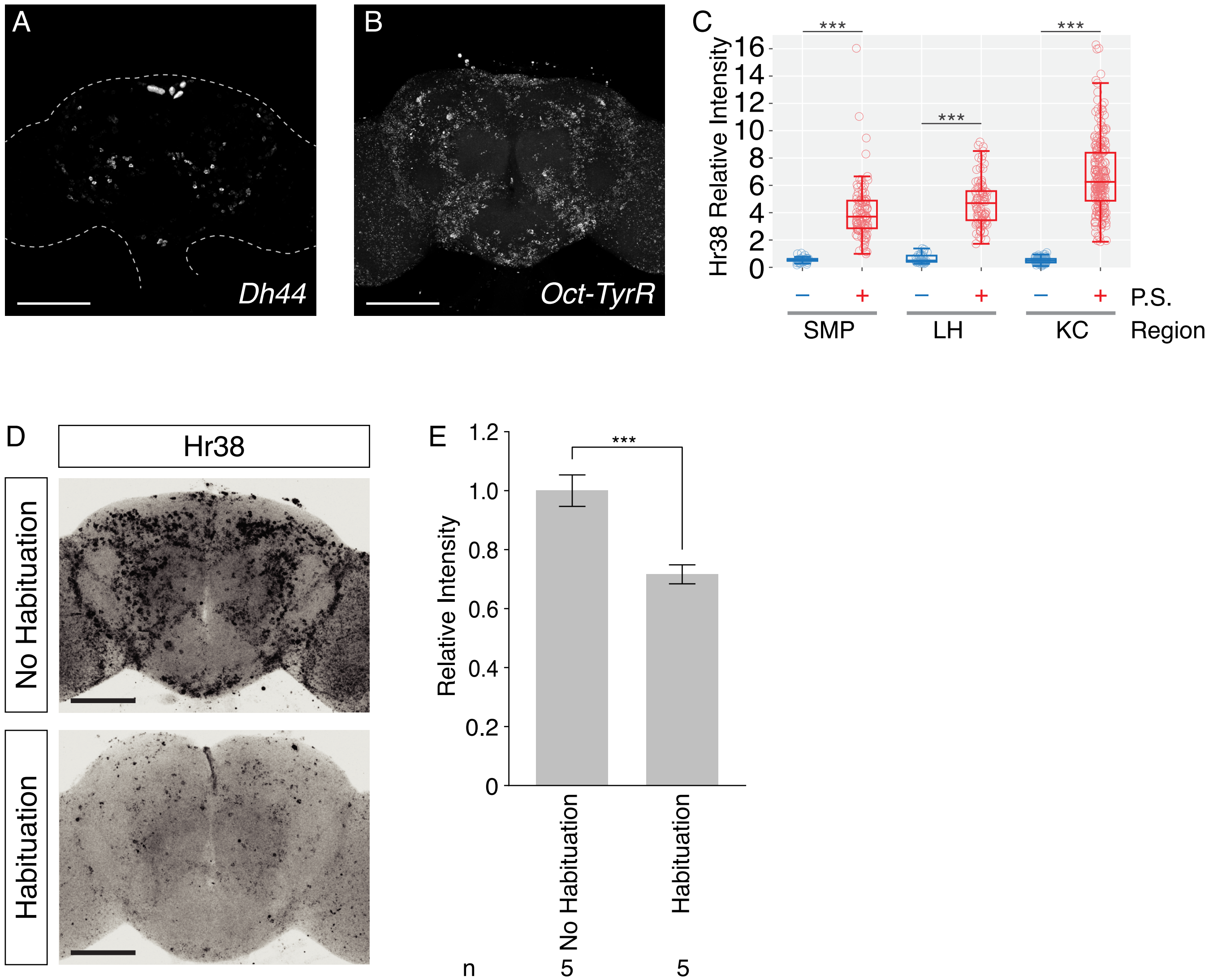

### Figure 2 - Supplement

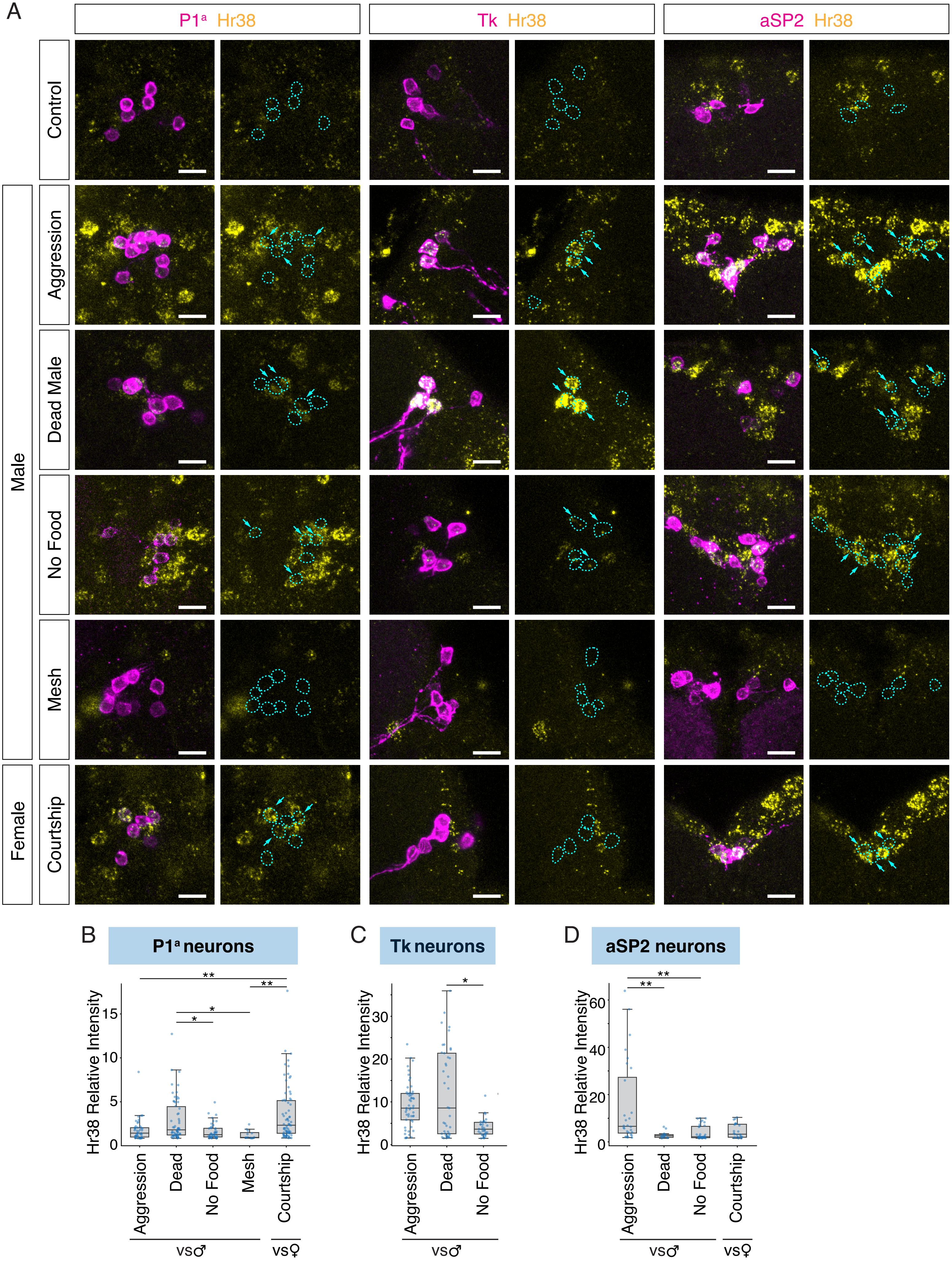

### Figure 3 - Supplement

Figure 3 - figure supplement

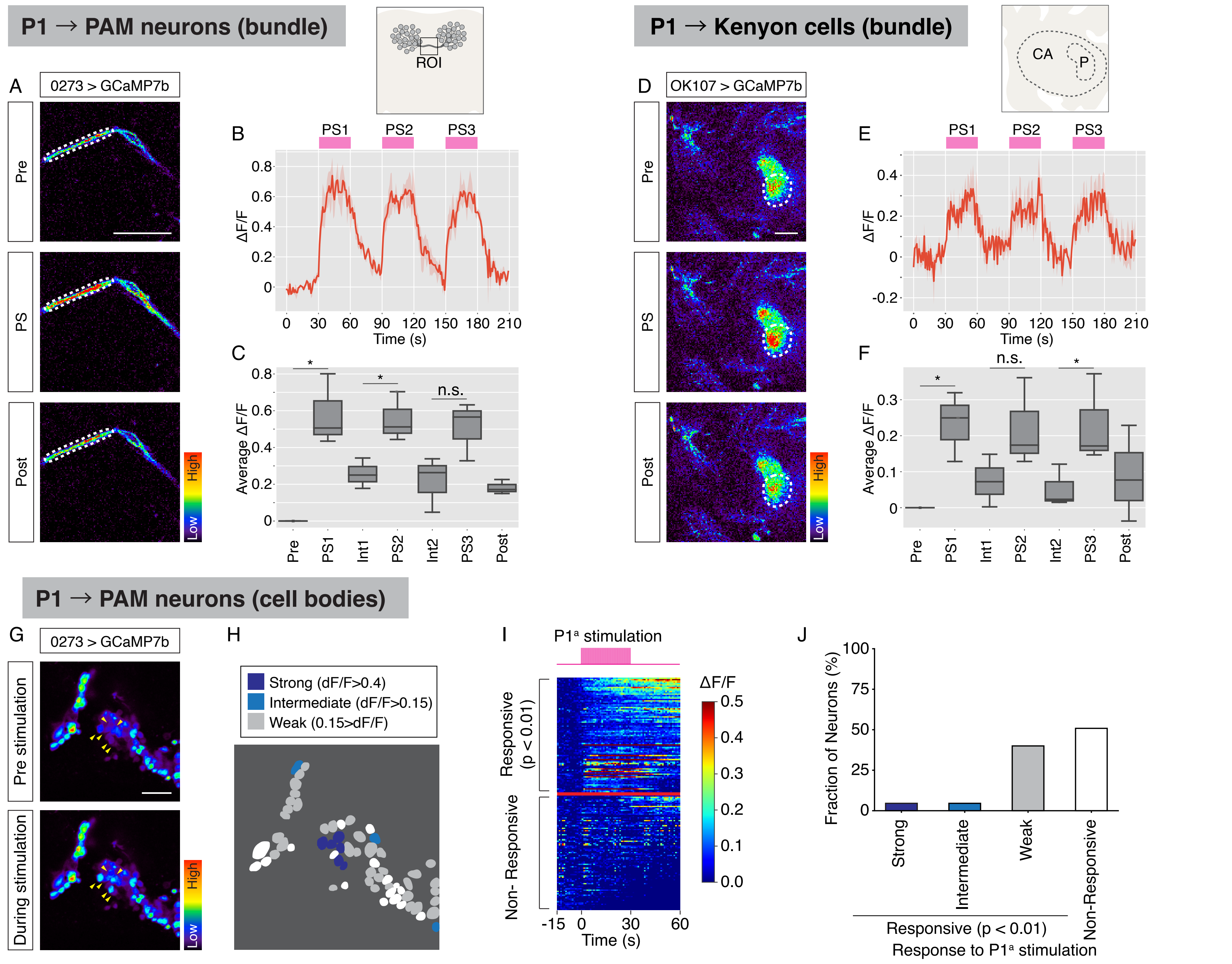

### Figure 4 - Supplement

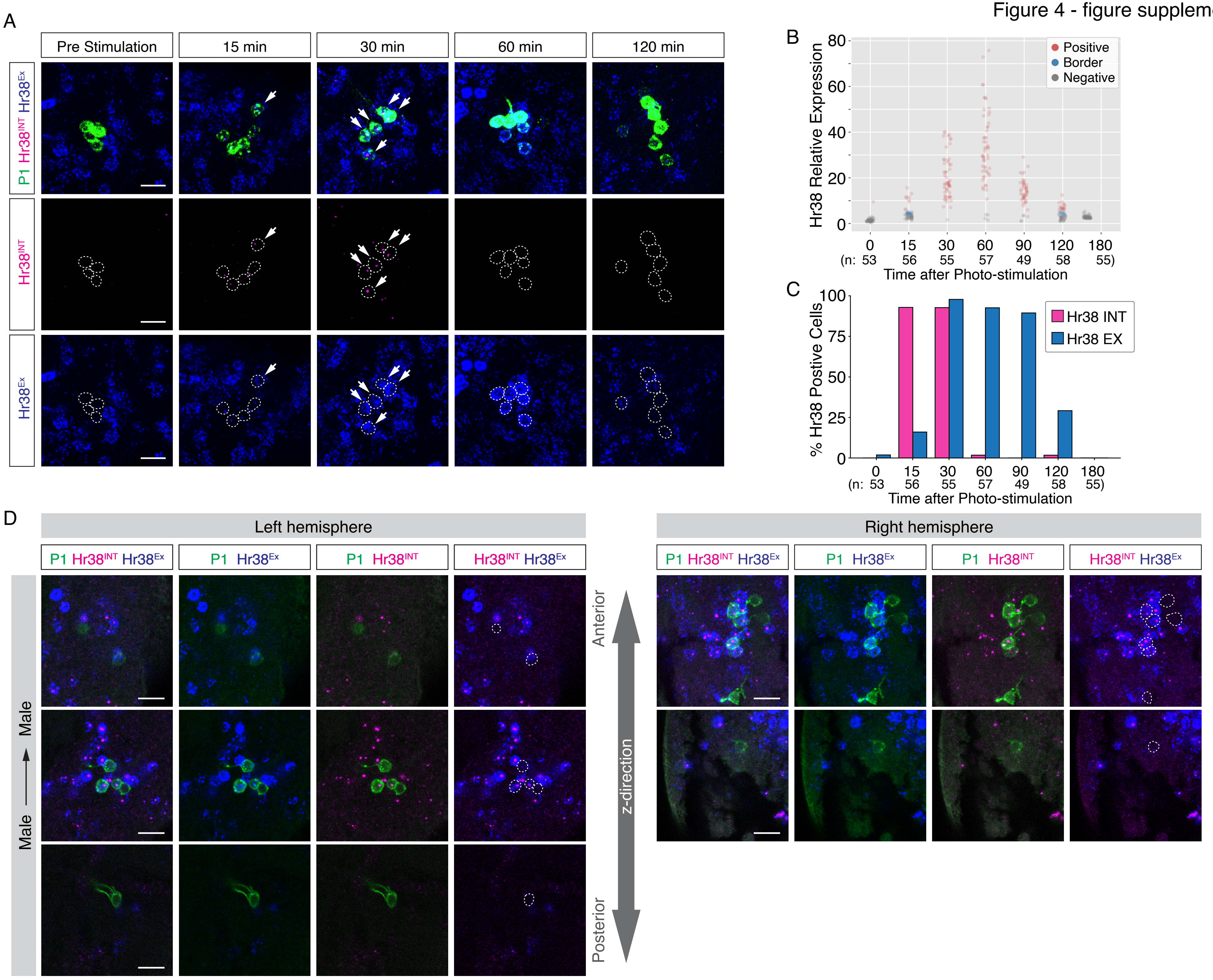
