## Supplementary Protocol for "HI-FISH: WHOLE BRAIN IN SITU MAPPING OF NEURONAL ACTIVATION IN DROSOPHILA DURING SOCIAL BEHAVIORS AND OPTOGENETIC STIMULATION"

**HI-FISH with *Hr38* probe (from sample preparation)**

**Sample preparation and Fixation**

Reagents

Ethanol

PBS

fixative: 4% PFA / PBS (prepare on the day of the experiment)

1. Habituating flies in the arena for the experiments.

As mechanical stimulation during transfer induces *Hr38* expression, transfer flies to the arena for the experiments ~ 3 hours before the experiment. It is possible to extend this period to overnight, but in that case food should be included in the arena.

2. Perform photo-stimulation or behavioral assay to induce *Hr38* expression.

3. For simple *Hr38* detection with the exonic probe, wait ~ 30 mins before beginning fixation and dissection.

4. Prefix the samples.

(This procedure can be omitted if there is a single brain to be dissected).

For dissecting multiple brains, prefix the samples before dissection.

Collect individual flies, soak them in cold ethanol to remove the wax. Transfer them into cold PBS.

Remove proboscis and open head cuticle quickly.

Transfer it to the cold fixative in a dish on ice (using a multi-well glass dish). After completing this procedure for all individual flies, keep the dish with the fixative on the ice.

Transfer one fly to the dissection dish (another multi-well glass dish) filled with ice-cold PBS and keep the samples on ice.

5. Dissect fly brains as quickly as possible.

Transfer one fly to another dissection dish (multi-well glass dish) filled with PBS (room temperature).

Dissect a brain and transfer it to the tube with fixative on ice. Dissection procedure is performed under a dissection microscope at room temperature.

5. Fixation (rotating/shaking, at 4C for at least 2 hours ~ O/N)

**Immunostaining (if desired)**

(In cases where HCR3.0 in situ is combined with immunostaining, perform the immunostaining first under RNase free condition, post-fix with 4%PFA for 20 min and then proceed to the following 'Pre-treatment' procedure. Otherwise, proceed to 'Pre-treatment' after 'Sample preparation and fixation.')

Perform regular immunostaining with fluorescent labeled antibodies on fly brains. For immunostaining under RNase free condition, use nuclease-free reagents.

Followings are some examples of these reagents and their suppliers.

Nuclease-Free Water	Thermo Fisher AM9932

Phosphate Buffered Saline (10x) pH7.4, RNase-free	Thermo Fisher AM9624

For reagents or buffers that are difficult to be prepared while maintaining RNase-free conditions (such as the antibody solution from DSHB hybridoma supernatant), RNasin Plus Ribonuclease Inhibitor is added (1/50 vol. Promega N2611).

**Pre-treatment**

Reagents (RNase free)

PBS

PBS 0.1% Triton X-100

PBS 0.5% Tween-20

2% acetic acid (ice-cold)

4% PFA

1. Wash samples 3 x 5 min with PBS / 0.1%Triton X-100 at room temperature to remove the fixative.
2. Permeabilization with PBS / 0.5% Tween-20 at room temperature for 20 min.
3. Rinse with PBS x 2.
4. Treat samples with ice-cold 2% acetic acid for 1 min (*).
5. Rinse with PBS x 2.
6. Post-fix with 4%PFA for 20 min on ice.
7. Wash samples 3 x 5 min with PBS / 0.1% Triton X-100 at room temperature.
8. Proceed to HCR v3.0 procedure.

* For most of applications targeting cytoplasmic mRNA detection, acid treatment can be omitted. For detecting nuclear signals using the intronic Hr38 probe, acid pre-treatment improves HCR signals (particularly, the signal of the Hr38 intron probe), but higher concentration/longer treatment decreases fluorescence signals (either native GFP fluorescence or Immunocytochemistry). Optimize the condition according to the type of labeling performed in each experiment. In general, 2% acetic acid for 1min is acceptable.

**HCR3.0**

Follow the protocol from Molecular Technologies with minor modifications

(https://files.molecularinstruments.com/MI-Protocol-RNAFISH-GenericSolution-Rev8.pdf)

Use the following hybridization buffer for better preservation of tissue morphology (simply replace Formamide in the original protocol with 8M Urea).

(ref. <https://doi.org/10.1016/j.ydbio.2017.11.015>)

30% Urea (8M ): final 2.4M

5x SSC

9mM citric acid (pH 6.0)

0.1% Tween20

50ug/ml heparin

1x Denhardt’s solution

10% dextran sulfate

Ordering information for modified hybridization buffer

8M Urea	Merck U4883

20x SSC	Thermo Fisher AM9763

Citric acid monohydrate	Merck C0706

50% Tween 20	Thermo Fisher 3005

Heparin sodium salt	Merck H3393

Denhard's Solution (50x)	Thermo Fisher 750018

Dextran Sulfate 50% Solution	Merck S4030

Hybridization reactions are performed in a 1.5 ml Eppendorf tube (nuclease-free).

~ 8 brains in ~ 50 µl hybridization solution (scaled down from the original protocol)

**Sample mounting**

I prefer to use Histodenz mounting solution, adopted from the Gradinaru lab procedure.

doi: 10.1016/j.cell.2014.07.017
